# Behavioral age-detection in individuals reconstructs minute-scale developmental transcriptomics

**DOI:** 10.1101/2025.11.01.686022

**Authors:** Nabeel S. Ganem, David Scher-Arazi, Sharon Inberg, Amit Zeisel, Shay Stern

## Abstract

Capturing rapid changes in gene-expression across development time is crucial for uncovering the molecular pathways that organize dynamic developmental processes. Here, we introduce ‘BehaveSeq’, a method that enables time reconstruction of minute-resolution developmental patterns of gene-expression by integrating longitudinal behavioral monitoring for precise age detection with genome-wide transcriptomic profiling in single individuals. By densely sampling individuals every ∼3 minutes during hours of *C. elegans* development, we uncovered thousands of genes exhibiting rapid and temporally structured expression trajectories. These genes were organized into distinct dynamical patterns associated with specific developmental functions. Moreover, time-reconstruction of transcriptional trajectories in serotonin-deficient individuals revealed time-specific neuromodulatory effects on developmental expression programs. Interestingly, using the same single-animal datasets, we were able to train a neural network model that accurately predicts the developmental age of each individual based on its individual-specific molecular signature. Overall, our method provides a new framework for revealing dynamic gene regulation with high temporal precision across developmental timescales.

## Introduction

Developmental processes are highly dynamic across time and are controlled by underlying changes in gene-expression states during these long timescales (Araya et al., 2014; Boeck et al., 2016; Hendriks et al., 2014; Large et al., 2025; Levin et al., 2012; Liu et al., 2022; Mody et al., 2001; Sun and Hobert, 2021; White et al., 1999). Thus, dissecting both slow and rapid changes in developmental patterns of gene expression is crucial for understanding the organization and mechanisms of developmental progression. Rapid changes in gene-expression patterns have been characterized in single genes using real-time fluorescent reporters (Chalfie et al., 1994; Jussila et al., 2022; Keil et al., 2017; Kinney et al., 2023; Pimmett et al., 2021). However, these measurements are limited to detecting one or a few genes simultaneously in a single experiment. On the other hand, measurements of temporal changes in global gene-expression states are usually based on averaging pools of individuals that are collected during discrete developmental windows following initial synchronization (Araya et al., 2014; Boeck et al., 2016; Levin et al., 2012; Meeuse et al., 2020; White et al., 2017). These methods may limit the temporal resolution of characterized gene-expression trajectories as single animals within a sampled population exhibit significant inter-individual variation in their developmental pace (Stern et al., 2017; Stojanovski et al., 2022; Irmler et al., 2004; Faerberg et al., 2021), even when born at the same time. Thus, a potential approach for significantly increasing the temporal resolution of developmental measurements of gene expression is to analyze genome-wide transcriptional profiles in single individuals for which the developmental age can be continuously and precisely defined. Then, by sorting these individual-specific gene expression profiles based on their developmental timing into a well-defined developmental trajectory, rapid gene expression changes over time can be revealed with high temporal resolution.

The development time of the nematode *C. elegans* is relatively short, with a total duration of 2.5 days from egg hatching to adulthood, and is composed of five developmental stages (L1 to L4 larval stages and the adult stage). During the transitions between developmental stages, *C. elegans* individuals show sleep-like behavioral states called lethargus, during which they become behaviorally inactive and molt (Cassada and Russell, 1975; Raizen et al., 2008; Stern et al., 2017). In previous studies, we developed and used a longitudinal multi-camera imaging system to study individual-specific behavioral changes across the full development time of *C. elegans* at high spatiotemporal resolution and under tightly controlled environmental conditions (Harel et al., 2024; Stern et al., 2017). In particular, this longitudinal behavioral imaging system allows inference of the exact developmental age of each single isolated individual, based on the individual-specific behavioral detection of the inactive lethargus periods exhibited during developmental transitions.

Here, we developed a new method (‘BehaveSeq’) that combines continuous behavioral age-detection and gene-expression profiling in the same single individuals, to reconstruct minute-resolution transcriptional trajectories during development. Gene-expression patterns revealed by this method include transcriptional states that change rapidly and show complex dynamics over developmental time, as well as genes with relatively gradual temporal changes in expression. Additionally, we found that specific gene groups identified by the reconstruction method to have similar temporal patterns, are enriched for unique developmental and cellular pathways. Furthermore, applying the newly developed method also to serotonin-deficient individuals revealed both shared and time-dependent changes in the developmental dynamics of gene-expression patterns affected by serotonin neuromodulation. Finally, the high-resolution measurements in single individuals enabled the development of a neural network model that predicts developmental age of an individual with minute-scale accuracy, based solely on its gene-expression profile. Overall, BehaveSeq integrates behavioral and molecular measurements in single animals for studying the temporal organization of gene-regulation with high temporal precision, at the individual and population level.

## Results

### Integration of behavioral age-detection and genome-wide transcriptional profiling in the same single individuals

We developed BehaveSeq, a new method that allows time-reconstruction of fast, minute-resolution temporal changes in gene expression states across development. In particular, the method combines long-term behavioral monitoring of isolated individuals for precise developmental age determination of each, and single-individual RNA sequencing in the same animals (Fig. 1A). The longitudinal behavioral monitoring was performed using a multi-camera imaging system that tracked the locomotory behavior of isolated wild-type *C. elegans* individuals (n=193) across their developmental trajectory, at high spatiotemporal resolution (3 fps) and under tightly controlled environmental conditions (Fig. 1A) (Harel et al., 2024; Stern et al., 2017). In this imaging system, individuals were grown in laser-cut multi-well plates filled with agar and a defined amount of UV-killed OP50 bacterial food to maintain homogeneous conditions across individuals during the experiment. Individuals growing in a food environment exhibit stereotyped long-term patterns of behavioral activity across development, divided by lethargus episodes of inactivity during molting, which robustly mark transitions between developmental stages (Fig. 1A; Fig. S1A) (Cassada and Russell, 1975; Stern et al., 2017). Different animals show variation in developmental timing, even when grown in the same environment and hatched at the same time (Fig. S1B,C) (Stern et al., 2017; Stojanovski et al., 2022; Faerberg et al., 2021). This natural developmental variation limits the temporal resolution when animals are pooled at specific developmental windows or sampling intervals using conventional sequencing methods, as each pool contains individuals at different development times within a time-range.

**Figure 1.**
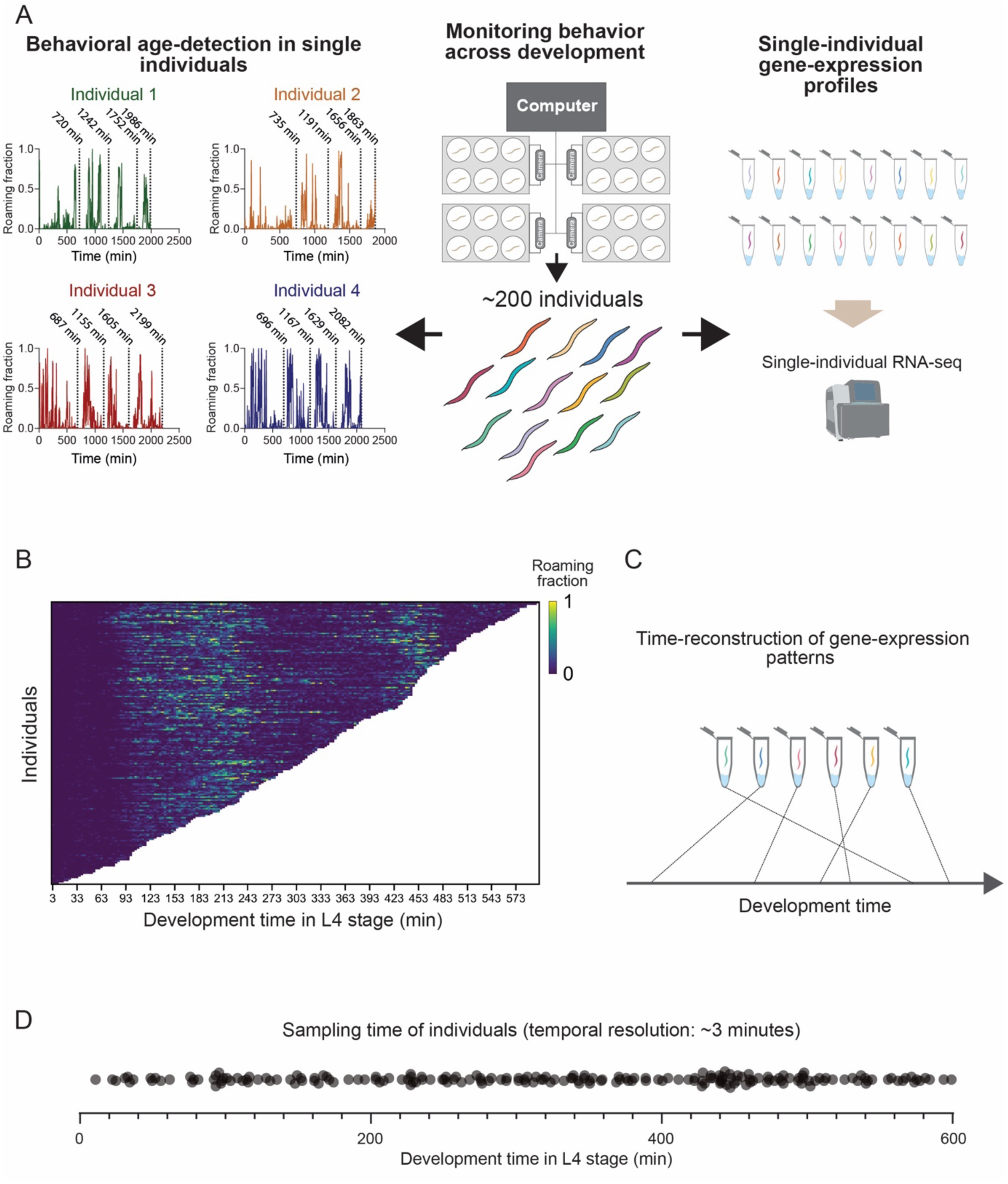
BehaveSeq methodology for combining single-animal measurements of behavioral and gene-expression states. **(A)** ‘BehaveSeq’ methodology overview. Middle: individual worms are monitored in isolation using a multi-camera imaging system to record locomotory behavior across all developmental stages at high spatiotemporal resolution and under controlled environmental conditions. Behavioral recordings of single individuals were terminated at different points throughout the L4 stage, followed by RNA sequencing in the same individuals, without losing their identity. Left: Automatic post-experiment behavioral quantification of single individuals measured in the longitudinal imaging system. Locomotion trajectories from raw imaging data (examples shown for 4 individuals) were analyzed to identify the developmental age of each individual and the exact times of transition across stages. For each individual, dashed black lines indicate 3 transitions between developmental stages and the time in which the individual was collected from the experiment (from left to right). Specific development times (min) are indicated above each dashed line. Right: single-animal RNA-seq libraries were prepared for each individual measured in the behavioral imaging system. Each library corresponds to a single individual at a defined developmental age. **(B)** Heatmap of behavioral roaming activity of wild-type individuals in the L4 stage (n=193), sorted by the developmental age of each individual within the L4 stage. Each row represents the behavioral trajectory of a single individual. Color indicates the fraction of time spent roaming in each of the 3 minute time-bins across the individual’s trajectory. **(C)** A schematics demonstrating the post-experiment sorting of individual RNA-seq samples in time, based on the behavioral age-detection of single animals. **(D)** Dot plot showing the time-distribution of individuals sampled within the L4 stage (n=193) worms. Each dot represents the developmental age of each single individual.

Here, we used the well-defined behavioral characterization to automatically and precisely identify, with high temporal resolution, the developmental age of each single individual in the population (Fig. 1A). Following the behavioral age-detection, we performed single-animal RNA sequencing using the Smart-seq2 protocol (Picelli et al., 2013), modified to measure RNA expression in single *C. elegans* individuals (Serra et al., 2018) (Fig. 1A). We sampled individuals with high temporal coverage during the L4 developmental stage by monitoring their behavior for approximately 28–39 hours post-hatching, covering the complete L1 to L3 stages and a variable duration in the L4 stage for each individual (Fig. 1B, Fig. S1D). These analyses resulted in densely sampled individuals at an average interval of 3.1 minutes across the 10 hours of the L4 stage, thus yielding high-resolution temporal snapshots of gene expression states during development (Fig. 1C,D).

In summary, we developed a new experimental framework that integrates behavioral age-detection and quantification of gene expression states in single individuals, enabling the reconstruction of high temporal-resolution transcriptional maps across development.

### Time-reconstruction of transcriptional dynamics using behavioral age-detection

Using the quantification of behavioral and gene-expression states in the same single individuals, we quantified the expression levels of 19,382 genes across 193 individuals sampled homogeneously throughout the L4 stage (Fig. 1D; Table S1). To verify that the time-reconstruction method based on behavioral age-detection in single animals recaptures underlying smooth patterns of gene-expression dynamics across development, we first tested whether individuals sampled close in time, even within a timescale of minutes, would show higher similarity in their gene-expression states. These analyses showed that individuals that were closer in developmental time since the start of the L4 stage (time since the L3-L4 transition) exhibited higher correlations between their gene-expression profiles (Fig. 2A). Specifically, samples taken from time-adjacent individuals separated by only a few minutes (≤5 min) showed significantly higher inter-individual similarity in gene-expression states (mean correlation = 0.92), compared to those separated by larger time gaps (mean correlation = 0.72) (Fig. 2B). In addition, a t-SNE map representing distances between gene-expression profiles across all sampled individuals showed a well-defined developmental trajectory, with individuals from adjacent timepoints clustering closer together (Fig. 2C). Importantly, we found that the reconstruction of temporal gene-expression patterns using the developmental time since the start of the L4 stage produced higher correlations among adjacent individuals, compared to time reconstruction using developmental time since hatching (mean correlation of 0.92 vs. 0.85) (Fig. 2A; Fig. S2A,B). Indeed, time reconstruction of transcriptional patterns of specific genes based on L4 initiation captured much finer features of temporal gene-expression dynamics over minutes (Fig. 2D), which also matched previously measured expression patterns by GFP reporters at high resolution (*lin-42* and *nhr-23*) (Kinney et al., 2023). The higher temporal accuracy obtained by taking into account time since the start of the L4 stage suggests that transitions across larval developmental stages serve as ‘milestones’ for initiating expression patterns (Levin et al., 2012), thereby resulting in more accurate time reconstruction in our setup.

**Figure 2.**
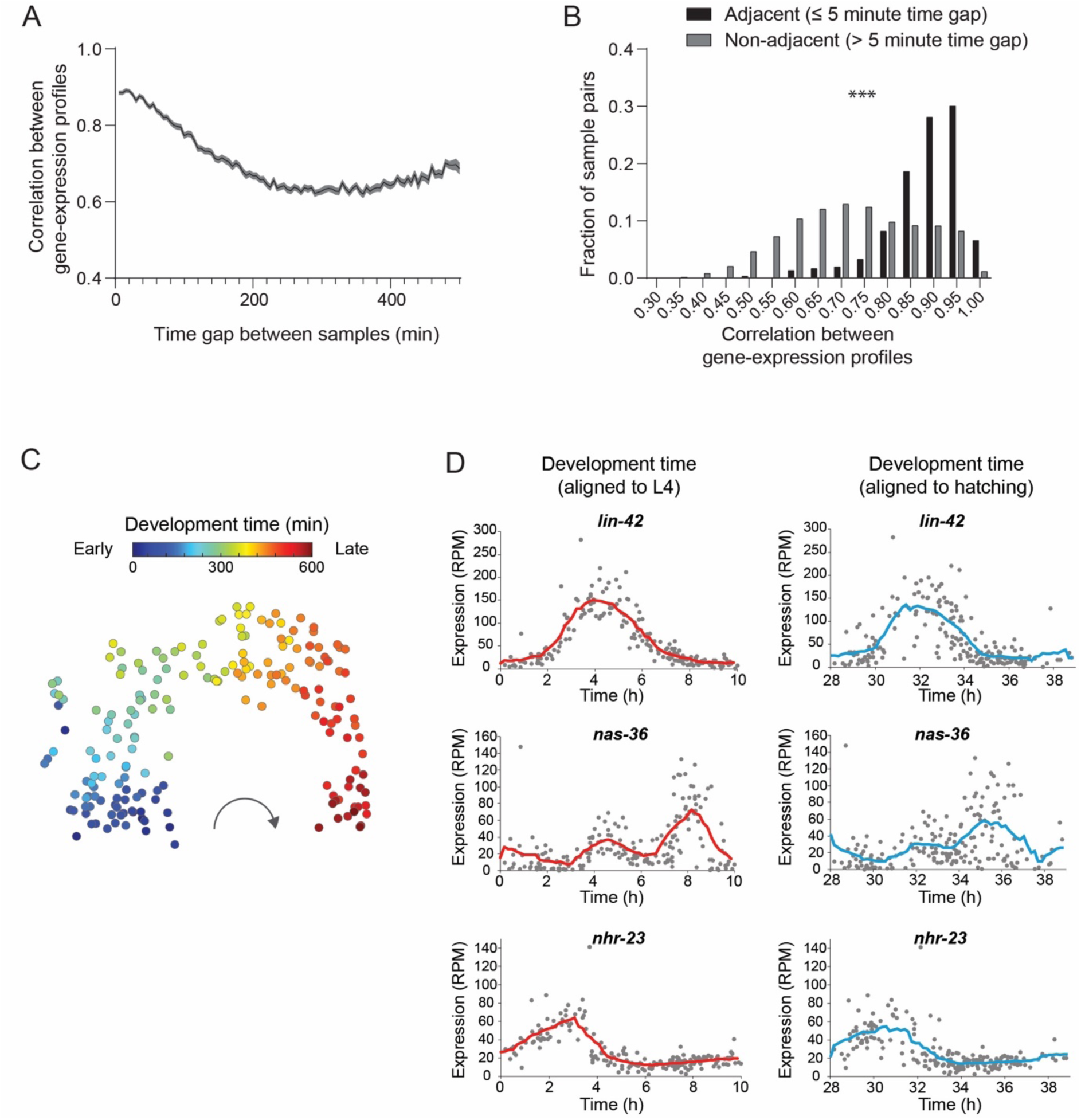
Reconstruction of high-resolution developmental patterns of gene-expression based on behavioral age-detection. **(A)** Average correlation between gene-expression states of pairs of wild-type individuals with a specific time gap between their developmental age. Shaded grey area represents standard error of the mean. Developmental age is aligned to the start of the L4 stage. **(B)** Comparison of distribution of correlation values between gene-expression states of pairs of wild-type individuals separated by less than 5 minutes in time (black) relative to pairs of individuals separated by more than 5 minutes (grey). *** P-value<0.001, Kolmogorov-Smirnov test. **(C)** t-distributed Stochastic Neighbor Embedding (t-SNE) visualization of all wild-type individual samples based on gene expression profiles. Each point represents the gene-expression profile of a single individual. Each single-individual sample is color-coded based on the behaviorally identified developmental age. **(D)** Temporal gene expression patterns of representative genes. Each dot represents the expression level (reads per million, RPM) of a specific gene in an individual’s RNA-seq sample, time-aligned based on developmental age since transition to the L4 stage (left, red) or since hatching (right, blue). Lines represent average gene-expression trajectories (smoothed using a window size of 1.45 hours, intervals of 0.17 hour).

Next, while our new method resolved temporal changes in gene-expression states on the scale of minutes, we sought to compare it with previous studies that used lower-resolution gene-expression measurements based on pooling individuals every hour during the experiment (Meeuse et al., 2020). To perform this comparison, we computationally reduced the resolution of our dataset by artificially pooling the expression levels of individuals within 1-hour windows since their hatching time (10 pooled samples from the original set of 193 individuals). We found that our artificially pooled gene-expression dataset closely matched the temporal expression patterns detected in previous studies where populations were pooled hourly (Fig. S2C,D), further demonstrating the efficiency of our method in reconstructing the population’s gene-expression patterns based on individual-specific measurements. These results show the precise time-reconstruction of the underlying dynamics of gene-expression via dense sampling of single individuals throughout the developmental trajectory.

### Reconstructed transcriptional trajectories reveal diverse and rapid temporal patterns of gene expression during development

The reconstruction of temporal patterns of gene expression uncovered a wide diversity of transcriptional trajectories during development. In particular, we found thousands of genes that showed both slow and fast developmental changes in gene expression. For example, among the genes that showed a decrease in expression during the developmental stage, we found *bath-28* and *ham-1*, which are expressed within neurons (Taylor et al., 2021) and showed continuous gene expression decay at different times within the stage (Fig. 3A). In addition, we found other genes that showed a non-homogeneous decrease in expression over time, composed of at least two episodes with different decay rates, such as *dre-1* and *nck-1*, which are involved in developmental timing and neuronal differentiation, respectively (Fielenbach et al., 2007; Horn et al., 2014; Mohamed and Chin-Sang, 2011) (Fig. 3A). Moreover, we identified genes that showed rapid oscillations within the developmental stage that were variable in structure, such as the *ugt-25*, *Y60A3A.23*, and *T16G1.2* genes that showed two peaks of activity, or the *pept-2* gene that showed faster oscillations of gene expression (Fig. 3B). The high sensitivity of the gene expression measurements over time also allowed us to identify genes that had rapid ON–OFF transitions from an inactive to an active state, and vice versa. For example, we found that *col-139* and *col-159*, which are involved in collagen synthesis during the molting cycle, as well as the *nspb-8* gene, which is predicted to encode a peptide, all showed transitions to ON or OFF states during specific development times that were variable among these genes (Fig. 3C).

**Figure 3.**
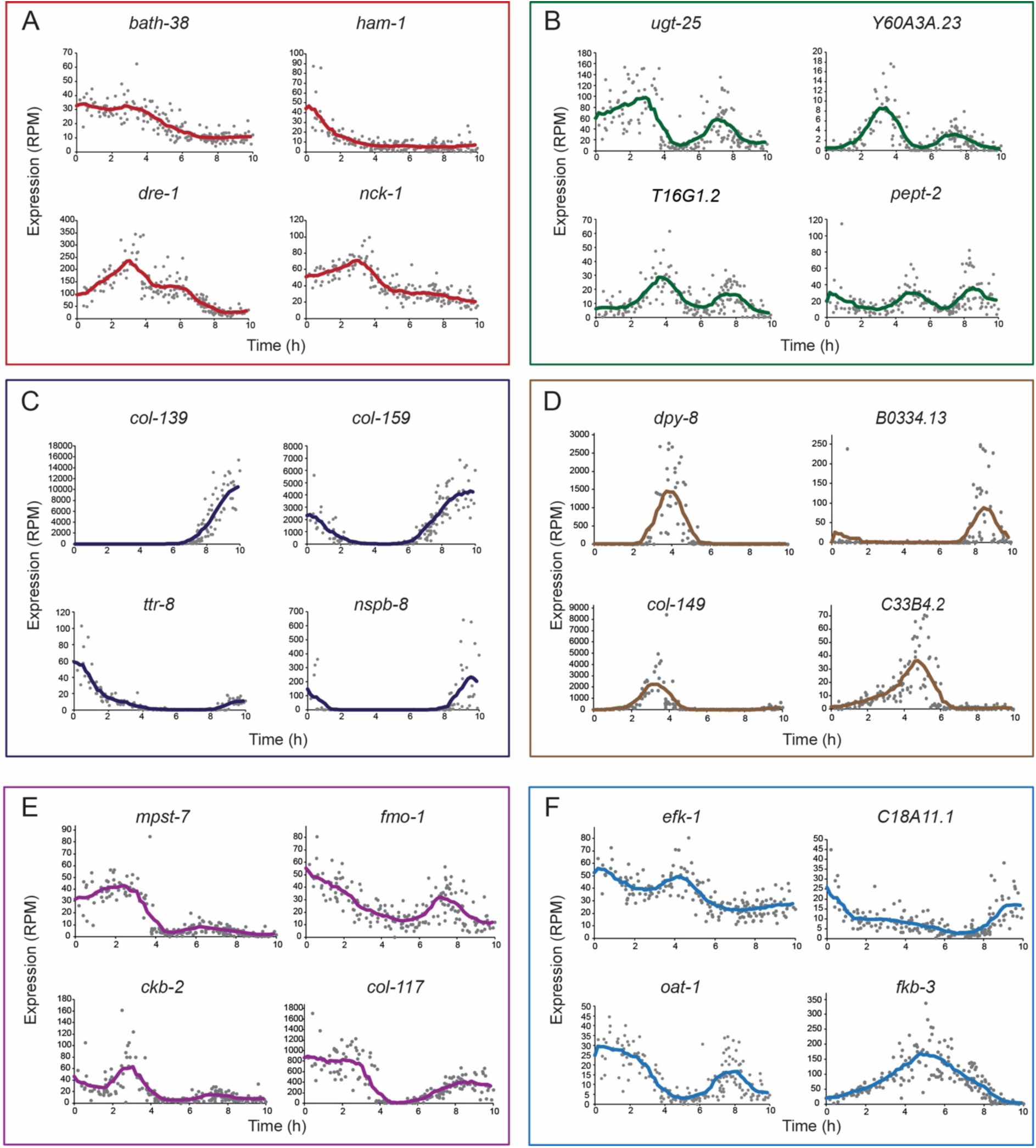
Detection of dynamic patterns of gene expression at minute-resolution during development. **(A–F)** Examples of the wide diversity of rapid patterns of gene-expression changes over development, uncovered by ‘BehaveSeq’. Each dot represents the expression level (RPM) of a specific gene in an individual’s RNA-seq sample, time-aligned based on the developmental age since transition to the L4 stage. Lines represent average gene-expression trajectories (smoothed using a window size of 1.45 hours, intervals of 0.17 hour).

In addition, we identified multiple genes with localized expression bursts that are surrounded by periods of inactivity during the stage, such as in *dpy-8*, *B0334.13*, and *col-149* (Fig. 3D). Within this group of ‘bursting’ genes, we captured asymmetric gene patterns where the rise and decay from peaks of activity are different, such as in the *C33B4.2* gene (Fig. 3D). Furthermore, the analysis of individual genes also uncovered various other types of gene expression patterns that had complex dynamics over time, such as *mpst-7* and *ckb-2*, which had a large peak of activity followed by a minor but still detectable increase in expression (Fig. 3E), as well as *efk-1*, which oscillated throughout its decrease over time, and *fkb-3*, which showed a gradual increase until the middle of the stage, followed by a symmetric decrease (Fig. 3F). Interestingly, while in all of the groups presented some of the genes have known developmental or physiological functions, the function of a large fraction of the detected genes is unknown (Table S1). Overall, these genes represent only a small subset of the full diversity of fast gene-expression patterns detected by ‘BehaveSeq’, underlying the global organization of transcriptional states during development.

### Uncovering gene groups sharing temporal patterns of expression and their associations with specific developmental functions

In addition to analyzing the diversity of gene-expression patterns at the single gene level (Fig. 3), as a complementary approach, we sought to identify specific groups of genes that show similar temporal patterns of transcriptional change across development. In addition, we also tested whether genes that share similar temporal patterns detected by our method, tend to be associated with specific developmental or cellular functions. Clustering analysis performed based on the temporal gene-expression patterns of all active genes (n=9,831 genes, 20 clusters, see Methods) revealed highly defined groups of genes showing distinct developmental structures of gene-expression (Fig. 4A,B). For example, we found that large clusters of genes (clusters 5,10,19; n=980, 407, 620 genes, respectively) showed a gradual increase or decrease in expression during the developmental stage. We also uncovered gene clusters that shared more complex transcriptional dynamics over development time (Fig. 4A,B). For instance, cluster 3 (n=377) showed an increase in gene-expression until 5 hours into the L4 stage and then a decline to lower levels compared to the start of the stage. In contrast, cluster 20 (n=207) showed a decrease in gene-expression until 2 hours into the L4 stage and then an increase in expression until 7 hours into the stage. In addition, clusters 2 and 14 (n=364 and 315, respectively) showed oscillatory patterns of gene-expression within the stage. Moreover, other gene groups such as clusters 12 and 13 (n=320 and 231, respectively), had an activity peak during mid-stage that was different in shape and time to each other (Fig. 4A,B). Overall, the gene-expression clusters revealed by the newly developed method demonstrate that the temporal transcriptional patterns of thousands of genes are organized in a highly structured manner within the *C. elegans* developmental programs.

**Figure 4.**
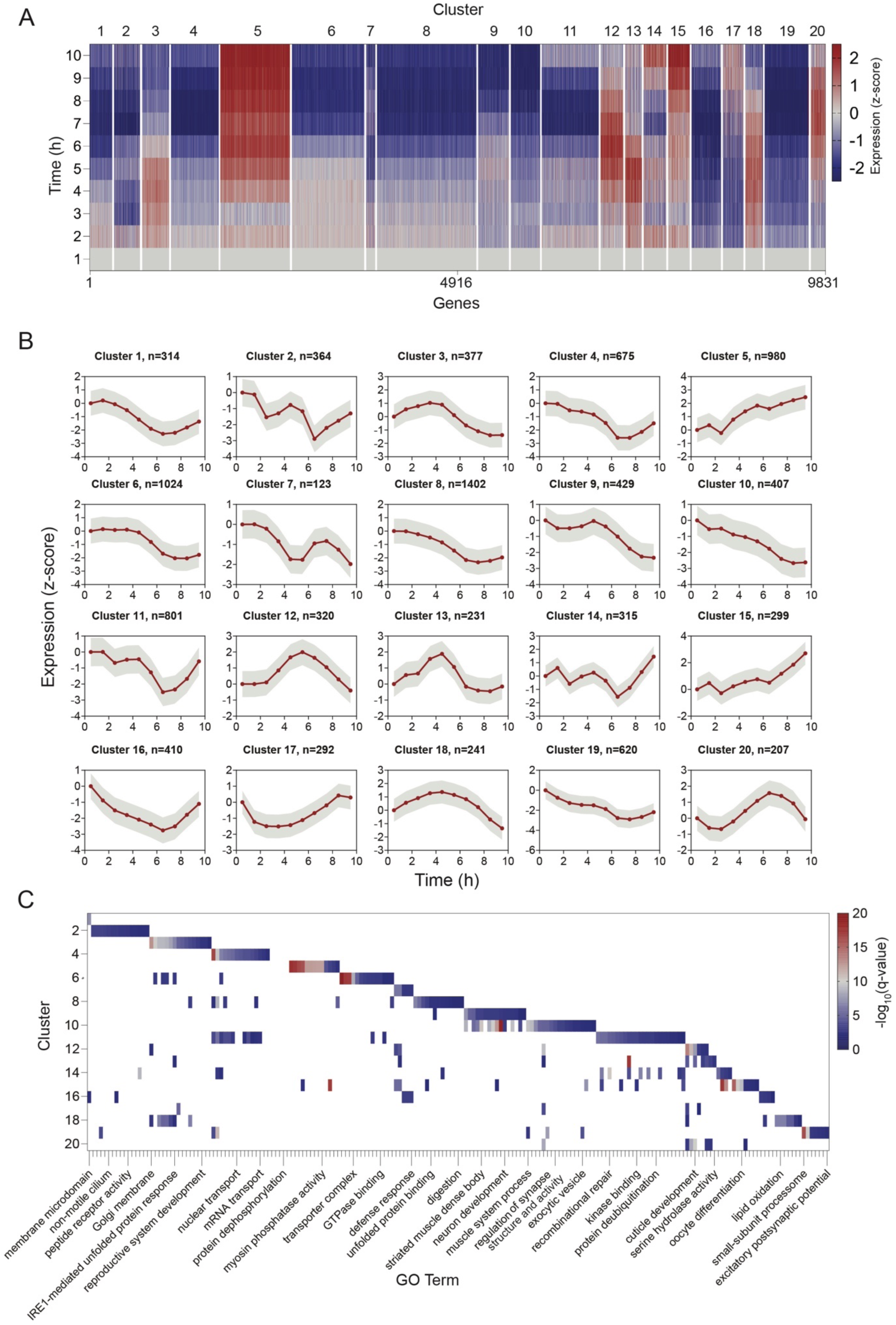
BehaveSeq reveals gene clusters associated with specific developmental pathways. **(A)** Heatmap of genes grouped by clusters (hierarchical clustering, see Methods) identified based on temporal gene-expression patterns (n=9831 active genes, see Methods). The x-axis represents individual genes ordered by cluster number, and the y-axis shows average expression of genes across 1-hour time-bins during the L4 stage. White lines separate clusters and the color code indicates normalized expression to first time bin (z-score). **(B)** Average temporal expression within each of the 20 gene clusters presented in (A) (smoothed using a window size of 1.5 hours, intervals of 0.49). Grey shaded area indicates the standard deviation. **(C)** Heatmap presents Gene Ontology (GO) enrichment analysis results for each gene cluster. Clusters are listed on the y-axis. Color code indicates the significance of enrichment (-log10(q-value)) for all significantly enriched GO biological terms (q-value<0.01). GO biological terms indicated by text in the figure represent only a partial list. Full list is in Supp Table S2.

To ask whether groups of genes identified in each cluster tend to be associated with specific developmental or cellular pathways, we further performed enrichment analysis for known biological functions (see Methods). We found that 15 of 20 clusters identified showed between 1 to 15 cluster-unique biological functions (q-value<0.01) (Fig. 4C; Table S2). These associated biological functions included specific neuronal and developmental processes, as well as molecular pathway. Specifically, genes within clusters 2 and 19, which exhibit either oscillatory or gradual decrease in gene expression over time, are significantly enriched for various peptide receptors expressed in the nervous system (Taylor et al., 2021). These genes include the guanylyl cyclases *gcy-9* and *gcy-13*, the acetylcholine receptors *acr-16* and *unc-29*, and the neuropeptide Y receptor *npr-12* (Table S2). Similarly, cluster 10, which also exhibits a gradual decline in expression during the stage, is enriched for genes known to function in neuronal development and synaptic transmission, such as *sax-7* and *unc-104*. Additionally, several clusters were enriched for genes that function in non-neuronal tissues and are involved in developmental processes, such as reproductive system development (cluster 3), cuticle development (clusters 12, 13, 17, and 18), and oocyte differentiation (cluster 15) (Table S2). Interestingly, other clusters were enriched for genes involved in chromatin organization and epigenetic regulation (Clusters 4 and 11), including *set-4* and *set-6*, and for specific molecular functions, such as the unfolded protein response (cluster 3) and DNA repair (clusters 4 and 11) (Fig. 4C; Table S2). In summary, the shared developmental and molecular functions identified within gene clusters uncover coordinated temporal programs of gene expression, underlying *C. elegans* development.

### Applying BehaveSeq across conditions reveals the neuromodulatory effect of serotonin on gene-expression patterns

To evaluate whether the temporal reconstruction of gene-expression patterns via behavioral age detection is robust across experimental conditions and can be utilized for comparison of transcriptional changes at specific developmental windows, we applied the ‘BehaveSeq’ framework to serotonin-deficient individuals, mutants for the TPH-1 serotonin-producing enzyme (Sze et al., 2000). Although multiple neuromodulatory pathways have been shown across species to affect neuronal and behavioral states (Bargmann and Marder, 2013; Marder, 2012), including in *C. elegans* (Horvitz et al., 1982; Flavell et al., 2013; Omura et al., 2012; Ripoll-Sánchez et al., 2023; Ali Nasser et al., 2023; Sanyal et al., 2004; de Bono and Bargmann, 1998), their impact on the temporal dynamics of gene expression during development is much less explored. We found that, similar to the wild-type population, integrating behavioral age detection with gene-expression states in *tph-1* individuals (n=164) enabled smooth reconstruction of transcriptional trajectories during development at a temporal resolution of ∼3.6 minutes (n=19,379 genes, see Methods) (Fig. 5A; Fig. S3A-C; Table S3). These high-resolution developmental profiles of gene expression in serotonin-deficient individuals revealed multiple gene clusters, composed of thousands of genes (n=9,952 total active genes, see Methods), showing highly variable patterns over developmental time (Fig. 5B; Fig. S3D,E; Table S4). To characterize specific time-dependent differences in gene-expression states caused by serotonin during development, we quantified for each gene its change in expression in all 1-hour time-bins between *tph-1* and wild-type individuals (Fig. 5C). We found that while many genes showed high temporal similarity between *tph-1* and wild-type individuals across developmental time (Fig. 5C,D), other genes showed consistent or time-dependent gene-expression changes over time. For instance, we found that while the *clec-47* and *T01D3.6* genes showed strong differences in the temporal patterns of expression across almost all 1-hour developmental windows in serotonin-deficient individuals, *sur-5*, *col-125* and *fat-5,* all showed significant changes that were limited to specific time-windows within the stage (Fig. 5C,E). More generally, we revealed 4,563 genes that showed significant change during at least one developmental window, 2023 genes that significantly changed during at least three developmental windows, and 441 genes that showed a more consistent change during half or more of the developmental windows (Fig. 5F, P-val<0.01). Interestingly, the group of genes that showed a significant difference during most of the developmental windows between *tph-1* and wild-type individuals (n = 441) was significantly enriched for genes involved in various metabolic and regulatory functions (Fig. 5G; Table S5). These functions include, for example, pathways involved in cellular respiration, nucleotide metabolism, tRNA activation, protein translation, unfolded protein binding, RNA localization, and regulation of gene-expression (Fig. 5G ; Table S5). These results suggest the effect of neuromodulation by serotonin on temporal patterns of genes, involved in core cellular and gene-regulatory functions across development. Moreover, we analyzed the cellular sites in which the genes that are temporally affected by serotonin operate. We found that serotonin tends to affect genes functioning in specific neuronal sites, including the NSM serotonergic neuron, which is one of the neuronal sites in which TPH-1 functions to synthesize serotonin (Horvitz et al., 1982; Dag et al., 2023; Zhang et al., 2005), as well as the PVD and PLM mechanosensory neurons (Chalfie and Sulston, 1981) and the AVK interneuron. In addition, genes temporally affected in *tph-1* individuals were also enriched in non-neuronal tissues such as the germline and pharynx (Fig. 5G ; Table S5). In summary, the reconstruction of developmental patterns of gene expression across conditions revealed both conserved and unique molecular patterns that are differentially affected over development time.

**Figure 5.**
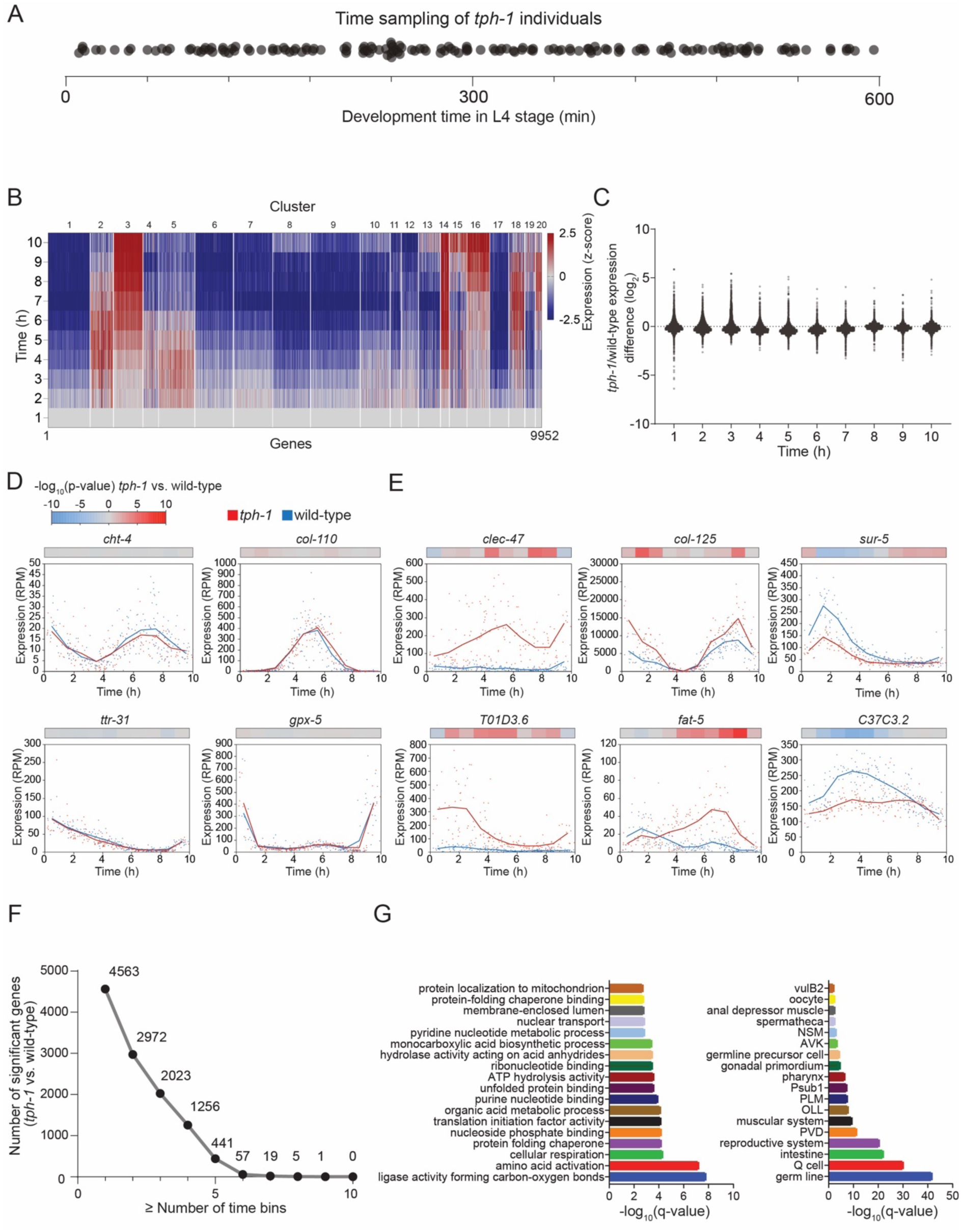
Developmental gene-expression dynamics in serotonin-deficient individuals. **(A)** Dot plot showing the time-distribution of *tph-1* serotonin-deficient individuals sampled within the L4 stage (n=164). Each dot represents the developmental age of each single individual within the stage. **(B)** Heatmap of genes grouped by clusters (hierarchical clustering, see Methods) identified based on temporal gene-expression patterns in *tph-1* individuals (n=9952 active genes, see Methods). The x-axis represents individual genes ordered by cluster number, and the y-axis shows average expression across 1-hour time-bins during the L4 stage. White lines separate clusters and color code indicates normalized expression to first time bin (z-score). **(C)** Gene-expression fold-change differences between serotonin-deficient and wild-type individuals at each 1-hour developmental time-bin (log2(*tph-1*/wild-type)) during the L4 stage. Each dot represents a single gene. **(D,E)** Gene-expression temporal profiles of representative genes showing either similar (D) or altered dynamics at specific developmental windows (E) between genotypes. Each dot represents the expression level of a gene in a single sample of N2 (blue) or *tph-1* (red) over time. Lines represent average expression levels (smoothed using a window size of 1.45 hours, intervals of 0.17 hours). Color bars above each panel indicate statistical significance of the difference in expression between genotypes in each of the 1-hour time bin. Mann-Whitney U test. P-values were corrected by FDR. **(F)** Plot indicates the number of genes that significantly changed across a specific number of developmental time bins or more, between *tph-1* and wild-type individuals. **(G)** Enrichment of biological functions (left) and tissue localization (right) within the group of genes that changes in at least half of the developmental time-bins between *tph-1* and wild-type (n=441). Shown are the 18 most significant results. Full list is in Supp Table S4.

### A neural network model for accurate prediction of an individual’s developmental age based on its transcriptional signature

Our results show that smooth transcriptional dynamics of gene expression can be accurately reconstructed by densely sampling individuals across the developmental trajectory, reflecting gene-expression states that are tightly locked to the time of development of each individual (Fig. 2). An open question is whether our individual-specific dataset can be further used for building a predictive model for precise identification of an individual’s age, based solely on its individual-specific gene-expression profile. We trained a multilayer perceptron neural network for regression by providing it with a training set of individual-specific transcriptional profiles and the associated developmental ages, quantified by the behavioral measurements (see Methods). We then tested the trained neural network by predicting the developmental age of a new set of individuals, based only on their gene-expression profiles (Fig. 6A). We found that by training a neural network using a set of 9,474 genes that showed activity over time across the conditions tested (see Methods), we generated a highly precise model that can predict developmental age with an average error of 18.7 minutes across trials (Fig. 6B,C). As expected, decreasing the number of genes used for training the neural network by selecting a random subset of genes (ranging from 5 to 8,000) led to an increased error in age prediction, up to an average error of 70.7 minutes when using only 5 random genes for the prediction (Fig. 6B,C). However, while the prediction error increased as the set of genes used for training was smaller, it remained highly significant relative to the prediction error generated by a shuffled dataset (average error across trials of 164.1 minutes, see Methods) (Fig. 6B,C). These results imply that the global gene-expression state of each single individual in the population carries sufficient information for developmental age prediction, and that this information is decreased by selecting a smaller, random set of genes.

**Figure 6.**
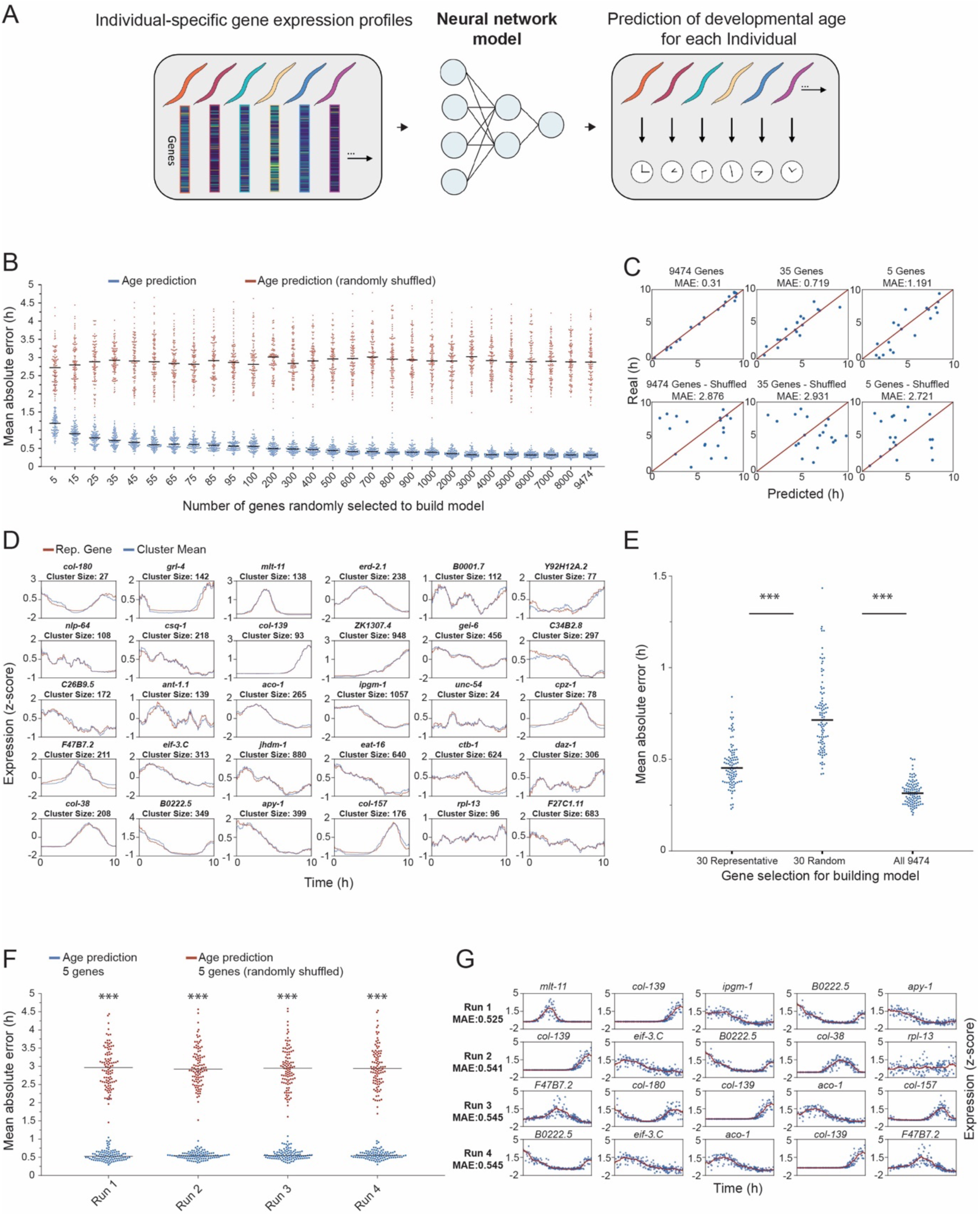
Neural network model for developmental age prediction in single individuals. **(A)** Schematics of a neural network model for predicting the developmental age of single individuals using individual-specific gene-expression profiles. **(B)** Shown is the prediction precision (average absolute error) of individuals’ developmental age by a neural network model generated by using random subsets of genes of different sizes (5 to total set of 9474 active genes, see Methods). For each of the gene subsets, shown are real prediction errors (blue) compared to prediction errors generated by a randomly shuffled dataset (red) (100 trials for each, see Methods). Each dot represents a single trial. Black lines mark the average precision across trials. **(C)** Examples of 3 trials from (B). For each trial shown are the observed (top) vs. random (bottom) predictions. Each dot represents the prediction of developmental age of a single individual vs. its true developmental age. **(D)** Representative genes (red) of distinct clusters (see Methods), identified as having the highest correlation (Pearson) to the mean expression of each cluster (blue). **(E)** The prediction precision of neural network models generated using the 30 representative genes as in (D), compared to a random selection of 30 genes, and to the total set of 9474 genes shared across conditions (100 trials). Each dot represents a single trial. Black lines mark the average precision across trials. ***P-value < 0.001, Mann-Whitney U Test. **(F,G)** Examples of 4 sets of 5 genes out of the 30 representative genes in (D). Shown are the average prediction errors of the sets of 5 genes with top prediction precision, relative to errors generated by randomly shuffled predictions (100 trials) (F) and the specific genes within these sets (G). ***P-value < 0.001, Mann-Whitney U Test.

To further test the possibility of minimizing the number of features selected, whilst still providing an accurate enough prediction, we sought to use a small non-random set of genes that contained a sufficient representation of the unique gene-expression dynamics present within the system. Using this set, the developmental age of an individual could be more efficiently located in time compared to a random selection of the same number of genes. We used 30 identified clusters of the gene-expression dynamics that were inferred within the overall set of genes measured (Fig. 6D), as these clusters represent groups of genes that are largely distinct in their gene-expression dynamics. In particular, for each of the identified clusters, we first found the mean expression pattern of the cluster, and then a representative gene within the cluster that showed the highest correlation (Pearson correlation) between its gene-expression pattern and the mean of the cluster, resulting in a set of 30 representative genes (Fig. 6D, see Methods). We then compared the prediction of developmental age using the non-random representative genes to the results of randomly selected genes. We found that neural networks trained using the 30 representative genes consistently showed significantly higher precision in predicting developmental age compared to the randomly selected set of 30 genes (average error across trials of 26.4 vs. 43.3 minutes) (Fig. 6E). Analyses using different numbers of clusters showed similar results (Fig. S4A). Next, to further minimize the number of representative genes required to build the model down to just 5, we randomly sampled (1,000 times) 5 genes out of the set of 30 representative genes. Out of these combinations, we found that the sets of genes that predicted with the highest precision reached an average error of 33 min, implying that it is possible to minimize the error in predicting age whilst only using 5 marker genes for the model (Fig. 6F,G).

To demonstrate the generality of our prediction method across conditions, we repeated our analyses to predict developmental age based on gene-expression states in serotonin-deficient *tph-1* individuals. Similar to wild-type animals, we were able to predict developmental age with high accuracy (average error 15.4 minutes) when using the total set of genes (n = 9474) (Fig. S4B). In addition, using sets of representative genes in the dataset of *tph-1* individuals, led to significantly higher precision of age prediction compared to random sets of genes (average error of 21.6 minutes) (Fig. S4C,D). Overall, the construction of a predictive neural network that produces consistent predictions provides further evidence for the tight association between the individual’s transcriptional signature and development time. Additionally, the identified small sets of marker genes with high predictive power could potentially be used to accurately determine the developmental age of each individual, without the need to measure their behavioral or morphological phenotypes.

## Discussion

A central challenge in understanding how developmental programs produce temporal changes in physiology, morphology, and behavior is to capture the underlying molecular states that rapidly change over the developmental trajectory. Traditional methods for measuring genome-wide gene expression changes over time are usually based on averaging expression levels during discrete developmental windows by pooling individuals within populations (Baugh et al., 2003; Boeck et al., 2016; Hashimshony et al., 2015; Levin et al., 2012; Meeuse et al., 2020; White et al., 2017), providing limited temporal resolution due to natural variability in developmental rate among these pooled individuals (Stern et al., 2017; Faerberg et al., 2021). Previous work showed that at the individual level, spontaneous behavior in *C. elegans* continuously evolves in a stereotyped manner across and within all developmental stages (Harel et al., 2024; Stern et al., 2017). In addition, during their development, individuals show behavioral quiescence periods (lethargus), marking the animal’s transitions across developmental stages (Cassada and Russell, 1975; Raizen et al., 2008; Stern et al., 2017). In this study, we leveraged the behavioral monitoring of individuals across development to introduce BehaveSeq, a new single-animal framework that uses behavioral dynamics to infer the precise developmental age of each single individual, followed by the quantification of the individual-specific transcriptional state using RNA-seq. The integration of these two measurements in the same single individuals enabled the time reconstruction of minute-resolution transcriptional trajectories from asynchronous individuals. We first demonstrated the newly developed method using a dense sampling of wild-type individuals during the fourth larval stage (L4). We revealed thousands of genes that showed rapid and non-homogenous expression changes over development time. In many cases, the fast changes in the expression of single genes, as well as their exact temporal structure, are hard to capture by using conventional methods which rely on pooling many individuals within much wider developmental windows. By performing clustering analysis based on the reconstructed temporal patterns of expression, we revealed coherent gene modules with distinct temporal profiles, reflecting shared regulation and functional coordination. Interestingly, many of these temporal clusters correspond to developmental processes, including nervous system development, cuticle remodeling, and reproduction, as well as to molecular pathways that change over development time, such as various peptide receptors and genes involved in epigenetic regulation.

To test how robust is the application of BehaveSeq across conditions, and to identify developmental differences in gene-expression trajectories, we also applied it to *tph-1* mutants, which are deficient for serotonin production (Sze et al., 2000). In addition to multiple genes that showed similar developmental patterns of gene expression in both *tph-1* and wild-type individuals, we also found genes that showed significant alterations in expression during specific developmental windows within the stage. The highly time-specific differences uncovered in serotonin-deficient individuals demonstrate the sensitivity of our method in detecting changes in expressions at specific developmental windows. In *C. elegans,* serotonin was shown to regulate behavioral activity across all developmental stages (Flavell et al., 2013; Stern et al., 2017). It is possible that in addition to the immediate effect of serotonin on neuronal activity via serotonin receptors (Dag et al., 2023), behavioral modifications in response to serotonin may also be encoded at the level of gene-expression states. Exploring the effects of genes that act downstream of serotonin would be valuable for understanding how neuromodulation affect behavior and physiology via changes in gene-expression. More generally, these results emphasize the utilization of BehaveSeq for comparison across conditions, which can be further used for studying the effects of various mutations and environmental perturbations on developmental patterns of gene expression.

In addition to reconstructing the population’s high-resolution transcriptional trajectories, we leveraged the discovered tight coupling between gene expression and development time at the individual level, to train a neural network to predict an individual’s developmental age based solely on its transcriptional profile. Using a multilayer perceptron trained on the full gene expression profiles of single individuals and their corresponding developmental age, we achieved a highly precise prediction of the developmental age when the network was tested on a new set of individuals (error<20 minutes). Importantly, we further demonstrated that using an even smaller set of selected genes, representing diverse temporal expression dynamics, was sufficient to retain high prediction accuracy, outperforming randomly selected gene sets of the same size. This finding highlights the existence of compact transcriptional signatures that reliably encode developmental timing and can be further used in future studies for age detection. We showed that this prediction framework is robust across both wild-type and serotonin-deficient individuals, suggesting that single-individual gene expression states contain sufficient temporal information for accurate age estimation across conditions, given a dense sampling of individuals over time.

While BehaveSeq was developed and validated in *C. elegans*, the core principle of using behavior as a continuous developmental marker to infer high-resolution temporal trajectories of gene-expression can be applicable to other systems. Across many species, behavior follows structured developmental transitions (Godoy-Herrera et al., 1984; Kimmel et al., 1974; Sokolowski et al., 1984; Vance et al., 2009; Wegman et al., 2010). By aligning individual-specific gene expression measurements based on time-resolved behavioral data, it may be possible to extract intrinsic developmental time-coordinates where external synchronization is impractical or where traditional developmental staging lacks resolution. This approach could also enable high-resolution time alignment of other molecular, physiological, or imaging data measured across single individuals. Altogether, the new method offers a powerful and generalizable approach for resolving minute-scale dynamics of gene expression, shedding light on the temporal organization of rapid gene regulatory processes during development.

## Methods

### Growth conditions and strains

*C. elegans* individuals were cultured on NGM agar media with OP50 *E. coli* as a food source according to standard protocols (Brenner, 1974). For behavioral imaging, isogenic worm populations were subjected to bleaching to isolate eggs. Single eggs were then transferred, using 2 uL of M9 buffer, to a custom-made multi-well plate created by laser cutting. Each worm was grown in its own circular arena across development (diameter of 10 mm). Each behavioral arena contained NGM agar supplemented with a specific amount of concentrated OP50 bacteria (10 uL of 1.5 OD), that were UV killed immediately after seeding the plates to prevent bacterial growth during the experiment (Stern et al., 2017). Wild-type Bristol N2 and MT15434 *tph-1*(mg280) strains were used in this study.

### Imaging system for tracking single individuals across development

Custom-made multi-camera imaging systems were used for longitudinal behavioral monitoring, each composed of six 12 MP USB3 cameras (Flea3, FLIR) and 35-mm high-resolution objectives (Edmund Optics) mounted on optical construction rails (Thorlabs) (Harel et al., 2024; Stern et al., 2017). Each camera captured images from six wells, each containing an individual grown in isolation. Movies were captured at 3 frames per second with a spatial resolution of ∼9.5 µm. To ensure uniform illumination of the imaging plates, identical LED backlights (Metaphase Technologies) and polarization sheets were utilized. Environmental conditions during the experiment were tightly controlled within a custom-made environmental chamber. The temperature was controlled using a Peltier element (TE Technologies), maintaining fluctuations within the range of 22.5 ± 0.1°C, humidity levels were maintained at 50 ± 5% through a sterile water reservoir, and external illumination was blocked, leaving the internal LED backlights as the sole source of illumination. Movies were captured from the cameras using commercial software (FlyCapture, FLIR) and saved onto two servers, with three cameras connected to each server.

### Imaging data processing for extracting locomotion trajectories and development time

To analyze the behavioral trajectories of animals throughout the experiment, we utilized custom scripts written in MATLAB (Mathworks) as previously described (Stern et al., 2017; Ali Nasser et al., 2023). In each frame of the video and for each behavioral arena, the worm was automatically identified as a moving object through background subtraction, and its XY position (center of mass) was monitored. For each experiment, approximately 350,000–450,000 frames per individual were analyzed using ∼50 CPU cores in parallel. This allowed us to reconstruct the complete locomotory trajectory of individuals over days of measurements across development. The overall time spent on image processing ranged from 39 to 45 hours per experiment. The hatching time of each individual in the experiment was automatically determined by identifying the timepoint when activity became detectable in the behavioral arena and manually verified. The middle of the lethargus periods, during which animals cease locomotion and molt, are detected as the transition points between different stages of development (based on 10-second timescale speed, smoothed over 225 frames). For each individual, chronological development time in all developmental stages was calculated based on detected developmental transitions. The presented quantification of roaming and dwelling behavior as a measure of locomotory activity across 3-minute time-bins was performed as previously described (Ali Nasser et al., 2023; Stern et al., 2017). Behavioral data for all individuals were deposited in Mendeley Data - (https://data.mendeley.com/preview/vm55hy4r4m?a=ebd9fe41-0d9c-449e-a08b-571893f98de3).

### Single-individual transcriptome sequencing

Preparation of sequencing libraries was done according to established protocols (Picelli et al., 2013; Serra et al., 2018) with some modifications. In brief, following the behavioral experiment, individuals were pipetted in a volume of 2 μl of UltraPure water onto the side wall of a clear 0.2 ml PCR tube containing 2 μl of Lysis buffer. The PCR tube was gently placed on its side on the stage of a ZEISS Stemi 508 dissecting microscope, and the individual was cut into few fragments using a sterile 25 G 1.5-inch regular needle, spun down, shock-freezed in liquid nitrogen and stored at −80 °C until use. Library preparation for each single individual was according to the Illumina Nextera tagmentation protocol (Illumina), with DNA input and reagents adjusted for 20 ng of cDNA. The tagmented DNA was cleaned, and PCR amplification was performed. PCR primers were removed using Ampure XP beads, and the libraries were eluted and quantified using Qubit fluorometer and Qsep100 Bio-Fragment analyzer. Libraries were pooled and sequenced using the Illumina NextSeq (550/2000 systems). The obtained sequencing read fragments were aligned to the WBPS16 using the STAR aligner (Dobin et al., 2013) v2.7.9a. Expression levels of single gene were normalized in each sample by gene mapped counts per million total reads (RPM).

### Reconstruction of developmental patterns of gene expression

Genes with missing values or zero expression across all samples were excluded from the analysis. Following this initial filtering, our dataset contained 19382 and 19379 genes for the N2 and *tph-1* populations, respectively. To reconstruct temporal gene expression dynamics, we aligned each sample to its exact developmental age in minutes, based on the behavioral age-detection. For each gene, expression values were plotted along a unified developmental timeline. This produced continuous expression trajectories for all genes. To assess the robustness of association between gene-expression states and development time, we performed correlation analysis (Pearson) between each unique sample pair. Then, we split the pairs into two groups: those in close proximity (≤5 min) and those further apart in time (>5 min). In addition, we also analyzed correlations among pairs by splitting them into groups ranging from a time-gap of 5 to 600 minutes. We used t-distributed Stochastic Neighbor Embedding (t-SNE) to visualize the low-dimensional representation of the similarities between the individual gene-expression profiles. The embedding was generated by using MATLAB (*tsne* function, dimensions=2, perplexity=40, and learning rate=110).

### Clustering of gene expression profiles and enrichment analysis

For the clustering analyses we performed hierarchical clustering on a set of active genes, defined as having 7 reads in at least 30% of the samples (9831 and 9952 genes for the N2 and *tph-1* populations, respectively), using a pipeline built in Julia (v. 1.11.5). As a pre-processing stage, the average expression level of each gene was calculated across 1-hour time-bins, log2(n+1) transformed, and normalized to z-score. We then normalized the temporal expression profiles to the first time-bin. Distance between two genes was defined as follows:

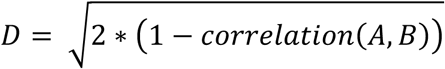

Where A and B are two different genes. The resulting distance matrix served as input to the *hclust* function of the *Clustering.jl* package with presquared ward linkage. Gene Ontology (GO) enrichment analysis was performed to identify molecular functions and cellular sites significantly enriched within gene clusters. The analysis was conducted using the WormBase enrichment analysis tool (https://wormbase.org/tools/enrichment/tea/tea.cgi), which utilize curated *C. elegans* specific annotations. Multiple testing correction was applied using the Benjamini-Hochberg false discovery rate (FDR) method. GO terms with an adjusted q-value < 0.01 were considered statistically significant.

### Statistical analysis

Pearson correlation coefficient (R) was calculated to assess the similarity between individual samples, based on L4 and total developmental time alignments. The Kolmogorov–Smirnov test was used to assess statistical significance of the difference in distributions of correlations among individual gene-expression profiles. To compare our N2 expression profiles with those of *Meeuse et al.* (2020), we first identified the intersection between our feature-selected genes (see Methods) and those with valid data in the reference dataset. To mimic the hourly pooled sampling of the reference, we binned samples at 1-hour intervals with a ±30-minute window. We then computed Pearson correlation for each shared gene, alongside a control in which gene expression patterns in the reference dataset were independently shuffled. Comparison focuses on developmental hours 28–36 in our dataset and hours 26–34 in *Meeuse et al.*, due to differences in growth temperature. An unpaired t-test was used to compare the correlations between our artificiality pooled dataset and Meeuse et al. against the correlations with shuffled gene expression profiles. For comparing expression patterns between N2 wild-type and *tph-1* individual in each 1-hour time bin, we used the Mann-Whitney U test (FDR corrected).

### Neural network model for age prediction based on individual-specific gene-expression profiles

Using the gene expression data of all active genes across the N2 and *tph-1* genotypes (n=9,474), a multilayer perceptron (MLP) with 2 hidden layers of 4 and 2 neurons, respectively, and an additional output layer of 1 neuron, was trained for regression to predict the time within the L4 stage that the sample was taken with a mean absolute error loss function (*trainnet*, MATLAB version R2025a). A lambda value of 7.3×10^-5^ was used. In each trial, for the N2 population of 193 samples, ∼90% (174) were used to train the network and ∼10% (19) were used to test the network. For the *tph-1* population of 164 samples, ∼90% (148) of samples were used to train and ∼10% (16) were used to test. To test network performance using a different number of features, we varied the number of genes used from 5 to 9474, at 28 unevenly spaced intervals. The genes were randomly selected without replacement in each of the 100 trials independently for each of the 28 selection numbers. In each trial, we also varied the training and test set used but kept the selection sequence the same across the feature numbers. We quantified the accuracy of each training by averaging the error across the test samples (mean absolute error). As a control, we shuffled the order of the predictions to calculate a random error in each training. For feature selection based on representative genes, we first clustered genes into 30 clusters using a pipeline built in MATLAB (R2025a). Before clustering, each gene was log_2_(n+1) transformed. Following that, genes were clustered via ward linkage with correlation metric using the *linkage* function. To find the representative gene of each cluster, the mean expression was calculated for each cluster and correlated with each gene (Pearson correlation) within that cluster. The gene with the highest correlation was identified as the representative gene of the cluster.

## Acknowledgments

We thank members of our laboratory for critical comments on the manuscript. We thank the Azrieli Technion Genomics Center. We thank WormBase, an online biological database for *C. elegans*, which is supported by Grant U41 HG002223 from the National Human Genome Research Institute at the NIH, the UK Medical Research Council, and the UK Biotechnology and Biological Sciences Research Council. Some strains used in this study were obtained from *Caenorhabditis* Genetics Center (CGC), which is funded by the NIH Office of Research Infrastructure Programs (P40 OD010440). NG acknowledges financial support from the Fein Family Postdoctoral Fellowship. DSA acknowledges financial support from the Ministry of Integration Doctoral Fellowship. SS acknowledges financial support from the European Research Council ERC-STG 851634 and Israel Science Foundation grant 3035/20.

---

**Figure S1.**
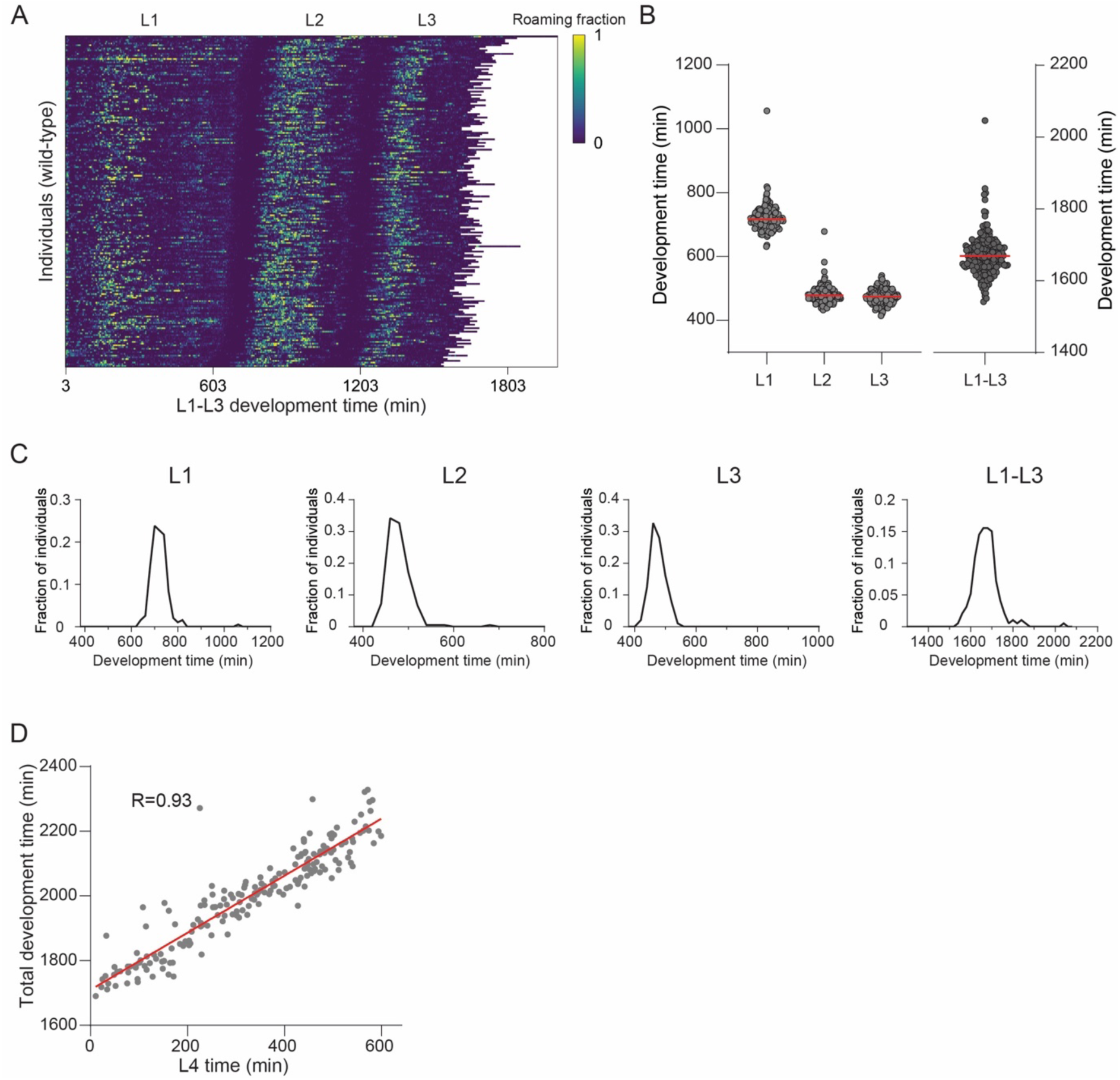
Inter-individual variation in development time. **(A)** Heatmap of roaming activity across the L1, L2, and L3 developmental stages in wild-type individuals (n = 193). Each row represents a single individual. Color indicates the fraction of time spent roaming in each of the 3-minute time-bins across the individual’s trajectory (see Methods). Heatmap is sorted based on L1 duration. **(B)** Plot shows the duration of each developmental stage and total time of the L1-L3 stages of wild-type individuals. Each dot represents a single individual. Red line marks the average duration. **(C)** Distributions of stage durations of each larval stage across individuals as in (B). **(D)** Correlation (Pearson) between the duration in the L4 stage and the total developmental time since hatching of wild-type individuals.

**Figure S2.**
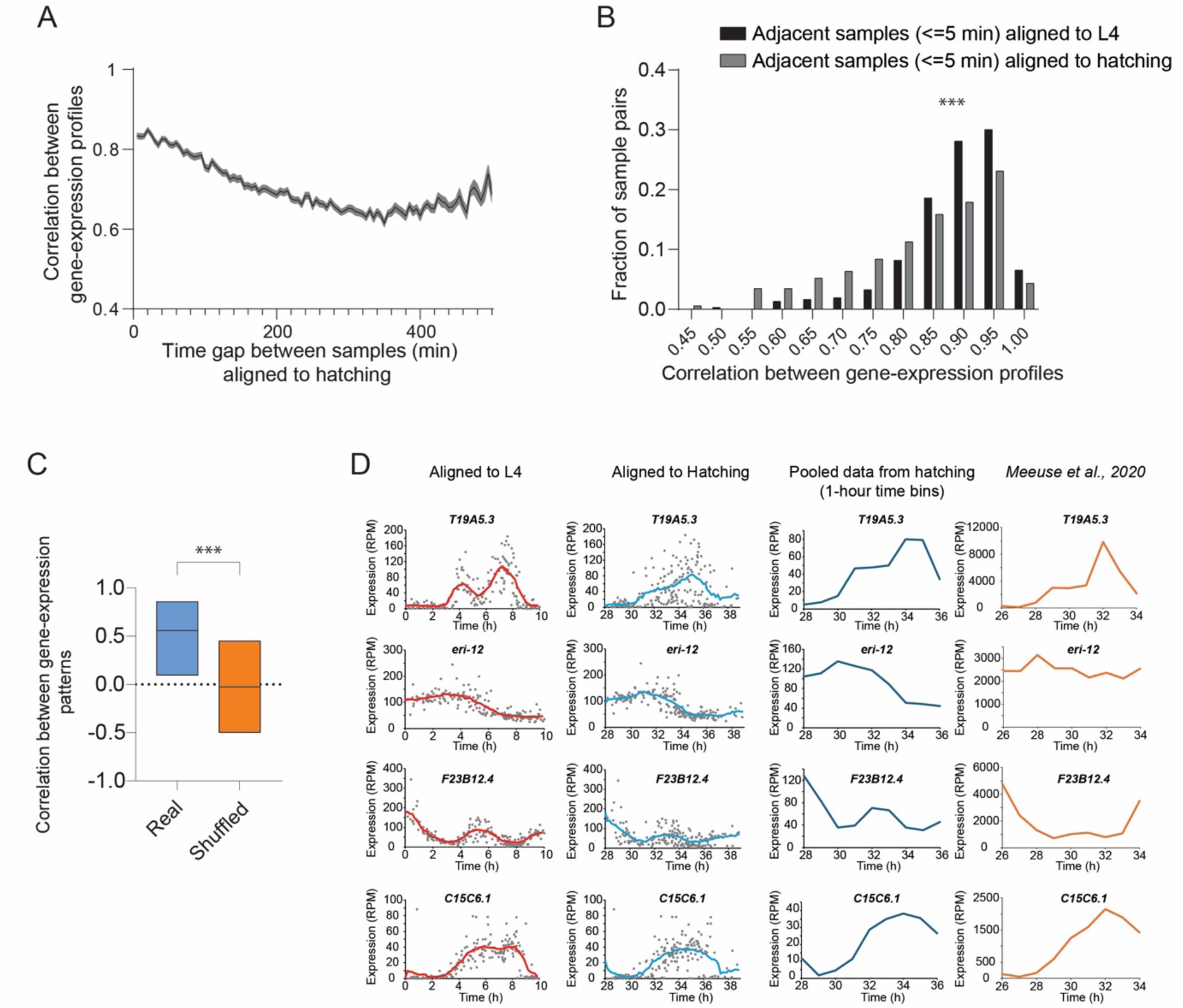
Similarity in transcriptional profiles of adjacent individuals and comparison to pooled dataset. **(A)** Average correlation between gene-expression states of pairs of individuals with a specific time gap between their developmental age. Shaded grey area represents standard error of the mean. Developmental age is aligned to the hatching time. **(B)** Comparison of distribution of correlation values between gene-expression states of pairs of individuals collected within 5 minutes of each other, based on L4 time-alignment (black) relative to time-alignment to hatching (grey). *** P-value<0.001, Kolmogorov-Smirnov test. **(C)** Correlation (Pearson) between temporal gene expression trajectories in our dataset, artificially pooled in each 1 hour time-bin (n=9740, see Methods) and a previously generated hourly pooled dataset (*Meeuse et al*. 2020) (Blue). Real correlations are compared to random correlations between shuffled datasets (orange). Boxes represent the variation (IQR) of the respective distributions. **** P-value<0.001, unpaired t-test. **(D)** Examples of gene expression profiles over time reconstructed from individuals aligned to L4 and hatching, as well as artificially pooled in each 1-hour time-bin in our dataset and in the hourly pooled dataset previously generated in *Meeuse et al*. (2020). Comparison focuses on developmental hours 28–36 in our dataset and hours 26–34 in *Meeuse et al.*, due to differences in growth temperature.

**Figure S3.**
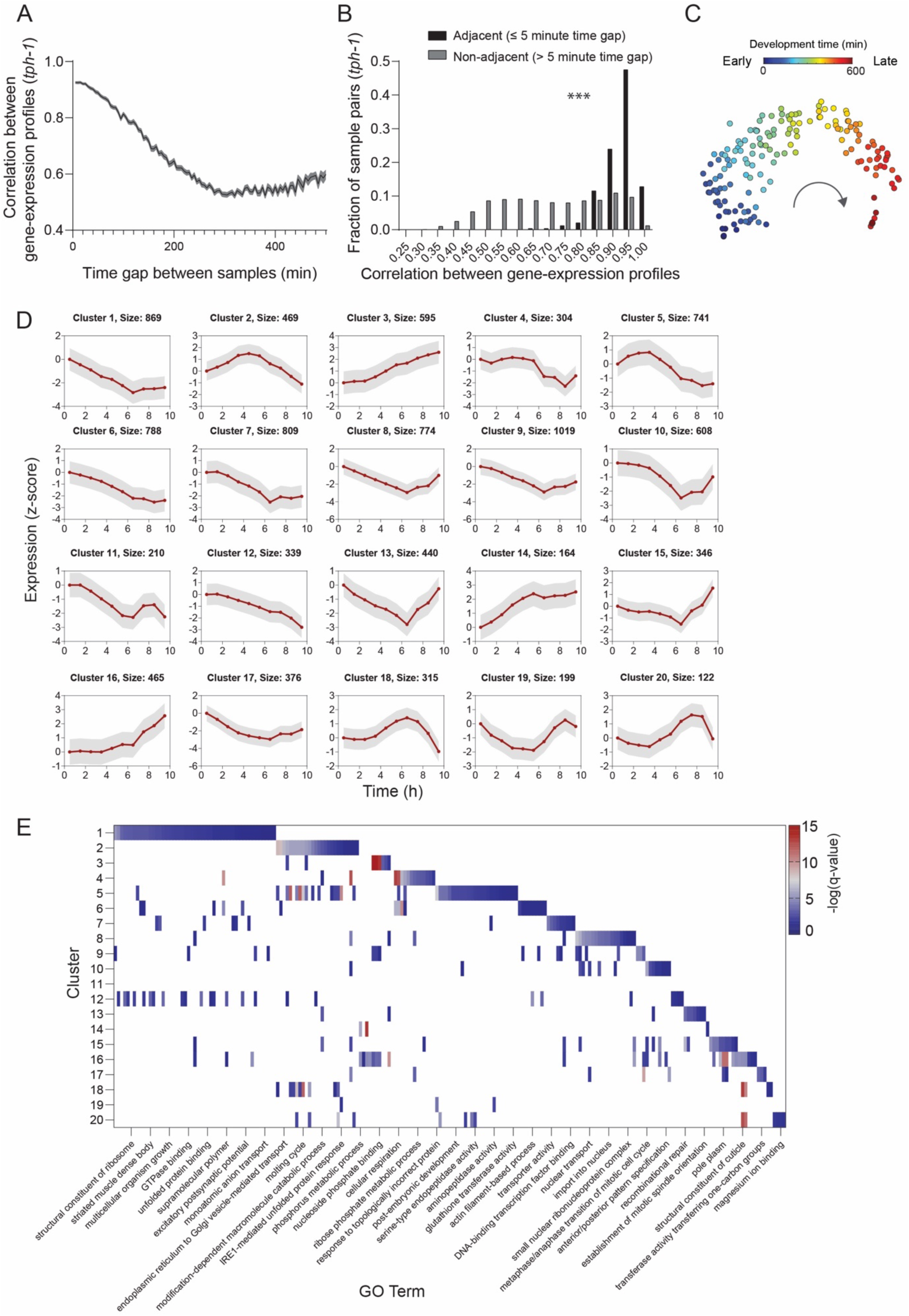
Time-reconstruction of temporal gene-expression patterns in *tph-1* serotonin-deficient individuals. **(A)** Average correlation between gene-expression states of pairs of *tph-1* individuals with a specific time gap between their developmental age. Shaded grey area represents standard error of the mean. Developmental age is aligned to the start of the L4 stage. **(B)** Comparison of distributions of correlation values between gene-expression states of pairs of *tph-1* individuals separated by less than or equal to 5 minutes in their developmental age (black) relative to pairs of individuals separated by more than 5 minutes (grey). *** P-value<0.001, Kolmogorov-Smirnov test. **(C)** t-distributed Stochastic Neighbor Embedding (t-SNE) visualization of all *tph-1* samples (n=164) based on gene expression profiles. Each point represents a single individual. Each single-individual sample is color-coded based on the identified developmental age. **(D)** Average temporal expression within each of the 20 gene clusters in Fig. 5B (smoothed using a window size of 1.5 hours, intervals of 0.49). Grey shaded area indicates the standard deviation. **(E)** Heatmap denotes Gene Ontology (GO) enrichment analysis results for each gene cluster generated from the *tph-1* dataset. Clusters are listed on the y-axis. Color code indicates the significance of enrichment for all significantly enriched GO biological functions (q-value<0.01). GO biological functions indicated by text in the figure represent only a partial list. Full list is in Supp Table S4.

**Figure S4.**
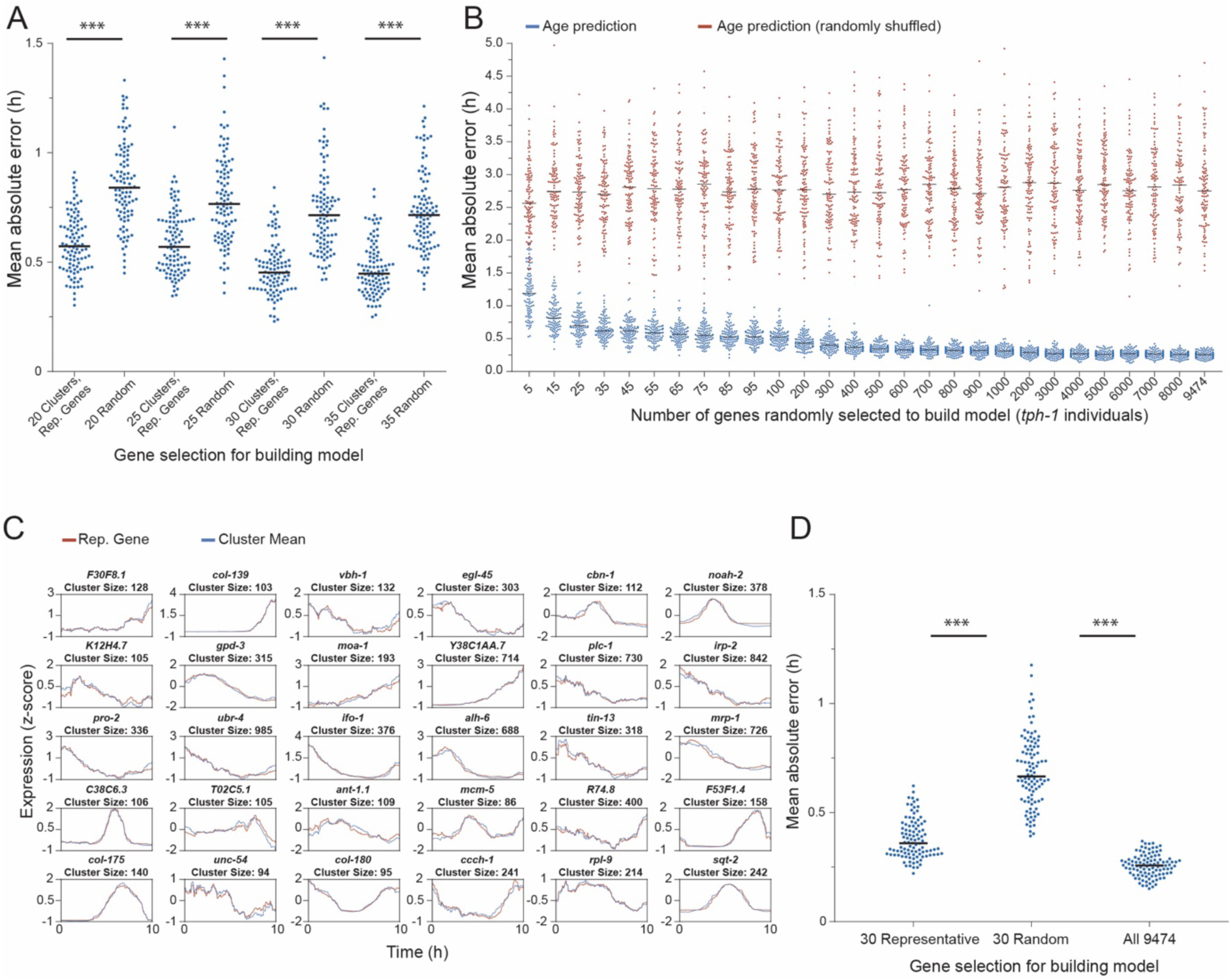
Developmental age prediction in wild-type and *tph-1* individuals. **(A)** Comparison of precision in developmental-age prediction using representative genes of wild-type individuals that correspond to variable number of generated clusters. Each dot represents a single trial. Black lines mark the average precision across trials. *** P-value<0.001 (Mann Whitney U test) for the difference in average error, compared to predictions using a randomly shuffled dataset. **(B)** Shown is the prediction precision (average absolute error) of *tph-1* individuals developmental-age by a neural network model generated using random subsets of genes of different sizes (5 to total set of 9474 active genes in both genotypes) (see Methods). **(C)** Representative genes (red) of distinct clusters (see Methods) generated from the *tph-1* individuals dataset. Each representative gene was identified as having the highest correlation (Pearson) to the mean expression of each cluster (blue). **(D)** The prediction precision of neural network models generated using the 30 representative genes as in (C), compared to a random selection of 30 genes, and to the total set of 9474 genes shared across genotypes (100 trials). Each dot represents a single trial. Black lines mark the average precision across trials. ***P-value < 0.001, Mann-Whitney U Test.

## Notes

### Competing Interest Statement

The authors have declared no competing interest.

